# Selenocyanate Derived Se-Incorporation into the Nitrogenase Fe Protein Cluster

**DOI:** 10.1101/2022.04.29.490034

**Authors:** Trixia M. Buscagan, Jens T. Kaiser, Douglas C. Rees

## Abstract

The nitrogenase Fe protein mediates ATP-dependent electron transfer to the nitrogenase MoFe protein during nitrogen fixation, in addition to catalyzing MoFe protein independent substrate (CO_2_) reduction and facilitating MoFe protein metallocluster biosynthesis. The precise role(s) of the Fe protein Fe_4_S_4_ cluster in some of these processes remains ill-defined. Herein, we report crystallographic data demonstrating ATP-dependent chalcogenide exchange at the Fe_4_S_4_ cluster of the nitrogenase Fe protein when potassium selenocyanate is used as the selenium source. The observed chalcogenide exchange illustrates that this Fe_4_S_4_ cluster is capable of core substitution reactions under certain conditions, adding to the Fe protein’s repertoire of unique properties.

## Introduction

The nitrogenase Fe protein has multiple roles, with its most famous role being ATP-dependent electron transfer to the MoFe protein during N_2_ fixation (Figure 1).^1–3^ The Fe protein also catalyzes MoFe protein-independent CO_2_-to-CO reduction,^4^ and participates in the biosynthesis of both the P-cluster and FeMo-cofactor.^5,6^ Unlike most Fe_4_S_4_ clusters in metalloproteins which adopt two oxidation states, the Fe protein cluster can span three oxidation states (2+/1+0).^7–9^ While both MgATP- and MgADP-binding to the Fe protein result in lower reduction potentials of the Fe_4_S_4_ cluster relative to the nucleotide-free state,^1^ only the MgATP bound state of the protein in the 1+ state is susceptible to rapid and complete iron chelation with bipyridine or bathophenanthroline.^10–13^ In the absence of nucleotide, iron chelation is slow, while MgADP inhibits chelation. Furthermore, the 2+ oxidized form of the Fe_4_S_4_ cluster undergoes ATP-dependent Fe chelation, yielding an intact Fe_2_S_2_ cluster.^13^ The origins of these unusual properties of the Fe-protein cluster are not well understood, but may reflect the solvent accessibility of the cluster and its positioning at the dimer interface.

**Figure 1.**
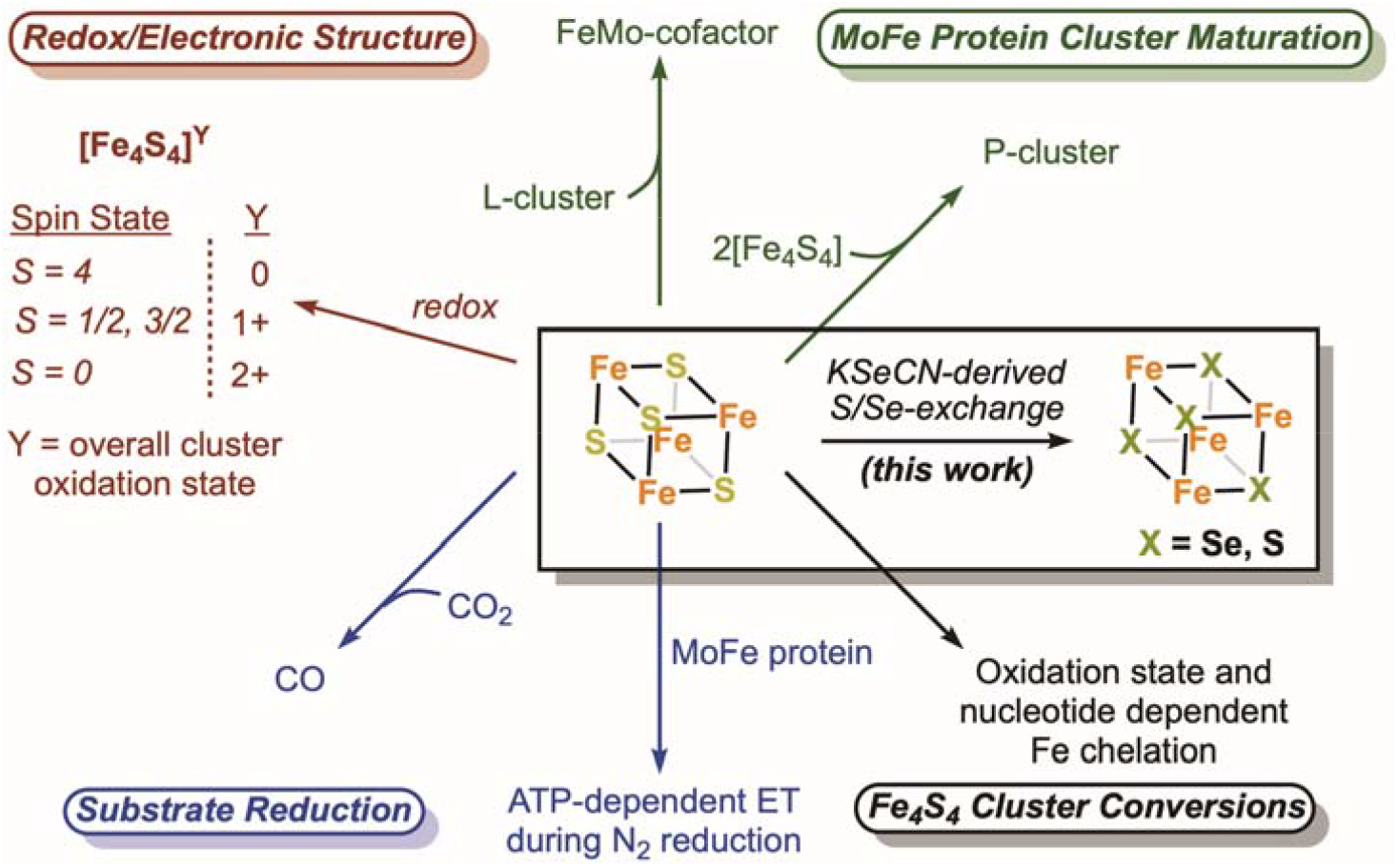
The nitrogenase Fe protein contains a Fe_4_S_4_ cluster with unique properties and participates in multiple reactions.

Our group has reported a crystallographic approach for quantifying selenium incorporation into the active site FeMo-cofactor of the MoFe protein.^14^ Key to this study was potassium selenocyanate (KSeCN), which like thiocyanate, is an alternative substrate for nitrogenase.^14,15^ Herein, using these conditions, we report a novel cluster conversion at the Fe protein in which the sulfide ligands of the Fe_4_S_4_ cluster exchange with “Se” from selenocyanate to yield an intact Fe_4_X_4_ cluster (X = Se, S) with Se-incorporation at all chalcogenide sites.

## Results

We initially observed Se-incorporation into the Fe protein cluster using the previously reported KSeCN turnover conditions developed for Se-incorporation at the FeMo-cofactor of the MoFe protein.^14^ Key components of the reaction include KSeCN as the selenium source, dithionite as the reductant, and an ATP regenerating system.^16^ Crystallization of the nitrogenase proteins from the concentrated reaction mixture was achieved by selecting conditions that are known to favor either MoFe protein or Fe protein crystals.^17,18^ The crystal structure at 1.51 Å resolution of the Se-incorporated Fe protein isolated from this reaction mixture is shown in Figure 2. The crystal form is isomorphous to the MgADP bound form of the Fe protein previously reported by our group,^17^ with the Fe protein molecular two-fold axis coincident with a crystallographic two-fold axis so that the asymmetric unit contains one subunit and half the cluster. The unique Fe1 and Fe2 sites are coordinated to Cys A97 and Cys A132, respectively, while the unique chalcogenide sites 3 and 4 are buried and surface exposed, respectively. The locations of the Se atoms within the protein structure were identified by collecting two sets of anomalous diffraction data: one above (12,668 eV) and one below (12,643 eV) the Se K-edge. Well-defined density was observed at both chalcogenide positions of the Fe_4_S_4_ cluster in the double difference anomalous Fourier map (Δanom_12668eV_-Δanom_12643eV_). Modeling the cluster exclusively as either the Fe_4_S_4_ or Fe_4_Se_4_ form resulted in substantial positive or negative difference density in the corresponding F_obs_-F_calc_ difference Fourier maps, respectively (Figure 2-figure supplement 1). Likewise, B-factors with lower or higher values at the core chalcogenide positions, relative to the iron cluster positions, were observed when the cluster was modeled exclusively as the all-sulfide vs all-selenide form, suggesting an under- vs over-modeling of electron density, respectively (Supplementary file 1). By fixing the chalcogenide B-factor values to a value similar to that of the Fe atoms, satisfactory mixed cluster models were obtained (see Methods for refinement details, Supplementary file 2, and Figure 2-figure supplement 2). The Se occupancies at the X3 and X4 positions are shown in Table 1, entry 1, with the X3 position exhibiting a greater extent of Se-incorporation relative to the X4 position.

**Table 1.** Summary of crystallographically determined Se occupancies for KSeCN derived Se-incorporation at the Fe protein cluster under various conditions. The occupancies for the X3 and X4 chalcogenide positions were determined in triplicate^§^ by analyzing three crystals prepared from a specified set of reaction conditions. For occupancy values corresponding to individual crystals (Supplementary file 3).

| Entry | Brief Description of reaction conditions* | X3 Occupancy<br>(Average + Standard Deviation) | X4 Occupancy<br>(Average + Standard Deviation) |
| --- | --- | --- | --- |
| 1 | 22 mM KSeCN, w/ MoFe protein | 0.51 ± 0.09 | 0.43 ± 0.06 |
| 2 | 22 mM KSeCN | 0.58 ± 0.03 | 0.38 ± 0.05 |
| 3 | 11 mM KSeCN | 0.07 ± 0.02 | 0.06 ± 0.03 |
| 4 | 1 mM KSeCN | 0.02 ± 0.01 | 0.02 ± 0.01 |

\*See Methods for full description
<sup>§</sup>With the exception of entry 2 for which four crystals were analyzed.

**Figure 2.**
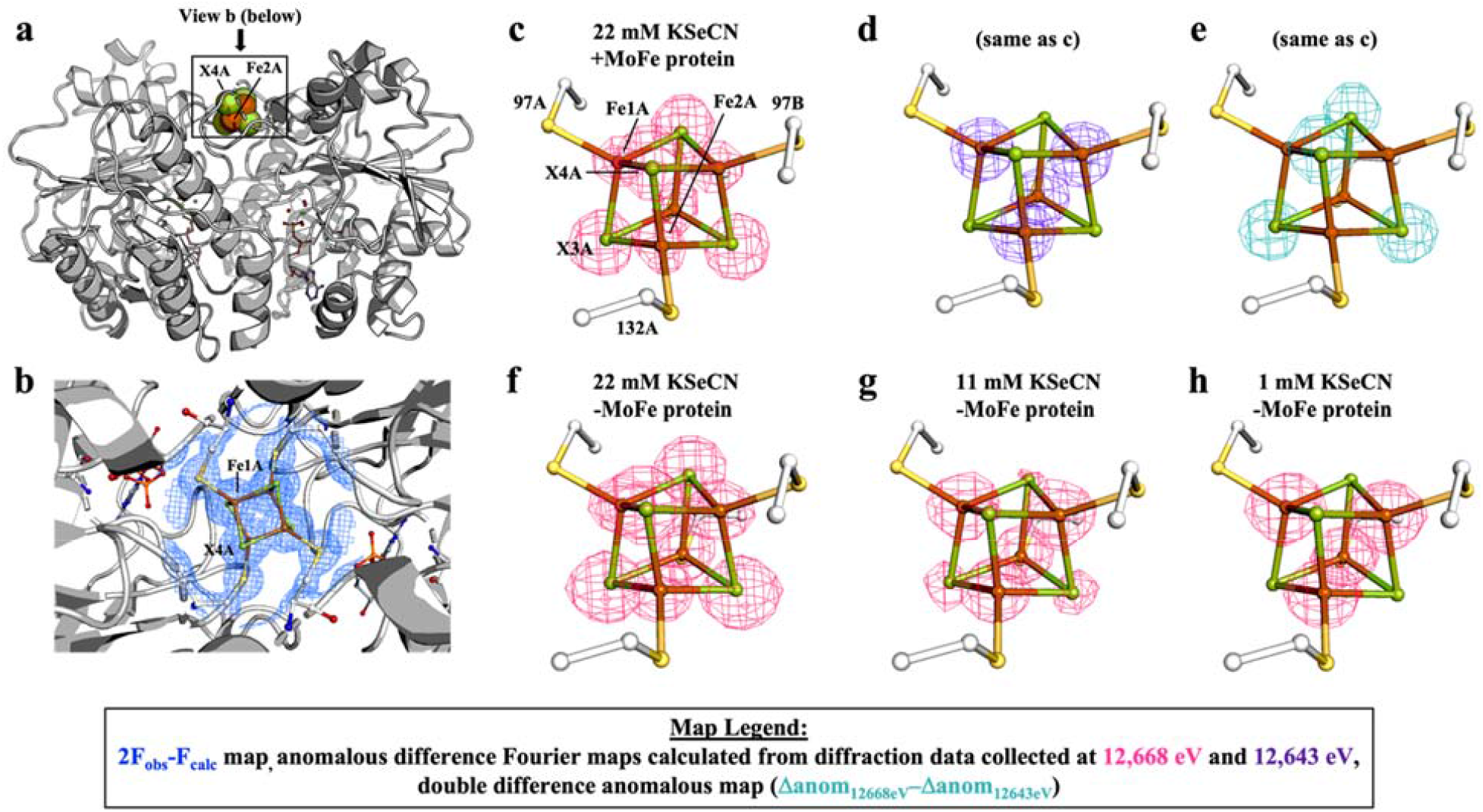
Pymol representation of the Se-incorporated Fe protein cluster at 1.51 Å resolution (PDB ID 7T4H), where the cluster chalcogenide (X) positions (green) feature a mixture of S and Se atoms. (a) Protein overview (b) with overlaid electron density (2F_obs_-F_calc_) map around the Fe_4_S_4_ cluster contoured at 1.5 σ (blue mesh) viewed with the dimer two-fold axis coincident with, and perpendicular to the plane of the paper, respectively. Anomalous difference Fourier maps calculated from diffraction data collected at (c) 12,668 eV contoured at 11.0 σ (magenta mesh, (d) 12,643 eV contoured at 11.0 σ (purple mesh), and (e) double difference (Δanom_12668eV_– Δanom_12643eV_) anomalous map contoured at 11.0 σ (teal mesh). (f-g) Anomalous difference Fourier maps calculated from diffraction data collected at 12,668 eV (magenta mesh) corresponding to crystals derived from reactions containing 22 (PDB ID 7TNE), 11 (PDB ID 7TPN), and 1 mM KSeCN (PDB ID 7TPO) contoured at 11.0, 7.0, and 5.0 σ, respectively. **Figure supplement 1.** Comparison of F_obs_-F_calc_ maps for cluster modeled as exclusively Fe_4_S_4_ vs Fe_4_Se_4_ forms. **Figure supplement 2.** Anomalous difference Fourier maps calculated from diffraction data collected at 12,668 eV for Se-free Fe protein crystals. **Figure supplement 3.** EPR spectrum of Fe protein used in control reaction with no MoFe protein. **Figure supplement 4.** Anomalous difference Fourier map calculated from diffraction data collected at 12,668 eV for ADP-bound Fe protein crystal soaked with KSeCN. **Figure supplement 5.** Double difference (Δanom_12668eV_– Δanom_12643eV_) anomalous map for Se-incorporated MoFe protein (FeMo-cofactor). **Figure supplement 6.** Fluorescence scan collected around Se K-edge energy for Se-incorporated Fe protein crystal. **Figure supplement 7.** EPR spectrum of purified Se-incorporated Fe protein.

To discern the essential components for Se-incorporation at the Fe protein cluster, control reactions were performed and the resultant Fe protein crystallized and subjected to X-ray diffraction. To determine whether the MoFe protein was required for Se-incorporation at the Fe protein cluster, the MoFe protein was omitted from the reaction (Table 1, entry 2). Surprisingly, Se-incorporation at the Fe protein cluster occurred in the absence of the MoFe protein as observed in the Δanom_12668eV_-Δanom_12643eV_ difference Fourier map. To rule out small amounts of contaminating MoFe protein, an EPR spectrum of the Fe protein used in the no MoFe protein control reaction was acquired (Figure 2-figure supplement 3); no signal corresponding to the *S* = 3/2 state of the FeMo-cofactor is observed. Additionally, the Fe protein used in the control was subjected to acetylene turnover conditions with no added MoFe protein. No ethylene formation was detected by gas chromatography, consistent with the absence of the MoFe protein. Performing the no MoFe protein reaction at lower KSeCN concentrations (11 and 1 mM KSeCN) resulted in a significant decrease in the intensities of the anomalous signals corresponding to the chalcogenide positions in the higher energy (12,668 eV) anomalous difference Fourier map, reflecting less Se-incorporation at the cluster (Figures 2e and 2f, and Table 1, entries 3 and 4). Having established that the MoFe protein is not required for Se-incorporation at the Fe protein, the nucleotide dependence of the reaction was examined. Omitting both the MoFe protein and ATP regeneration system from the reaction did not yield crystals suitable for X-ray diffraction (XRD) studies. To obtain suitable crystals for XRD, the control reaction was repeated, followed by addition of MgADP during the reaction work-up to form the MgADP bound form for crystallization. No Se-incorporation is observed in the anomalous difference Fourier map calculated from data collected at 12,668 eV, when the Fe protein and KSeCN are the sole components of the reaction (See Supplementary file 2, PDB ID 7TPY and Figure 2-figure supplement 2d). Additionally, when MgADP and KSeCN, but no MoFe protein or ATP regeneration system, are mixed with the Fe protein, no Se-incorporation at the Fe_4_S_4_ cluster occurs (Supplementary file 2, PDB ID 7TPZ and Figure 2-figure supplement 2e). Finally, in an attempt to observe a potential ligand-bound form of the Fe_4_S_4_ cluster, the MgADP bound crystal form was soaked with KSeCN; no density corresponding to ^-^SeCN, either near the Fe_4_S_4_ cluster or anywhere else in the protein structure, was observed (Figure 2-figure supplement 4).

## Discussion

The ability of iron-sulfur cluster containing metalloproteins to undergo a rich variety of cluster conversions and exchange reactions involving exogenous iron and sulfur species has been recognized since the pioneering work of Beinert.^19–22^ An orthogonal method for monitoring S-exchange in and out of clusters uses selenium as a structural surrogate of sulfur.^23,24^ Our group’s previous Se-incorporation results coupled with the results described herein highlight both the utility of this approach with nitrogenase and the selectivity of this process, under KSeCN turnover conditions. While the Fe protein cluster and the two-coordinate sulfides of the FeMo-cofactor undergo Se-incorporation, the P-cluster, which has been reported to undergo redox-dependent structural changes,^25,26^ has not yet been observed to undergo exchange of any of the constituent sulfides.

Relative to other iron-sulfur proteins, the Fe protein cluster exhibits unusual redox and electronic properties (Figure 1).^9,27,28^ Given the oxidation-state and nucleotide-dependent iron chelation and changes to spectroscopic features observed in the presence of MgATP,^27^ it has been proposed that MgATP-binding results in a conformational change that renders the cluster more accessible to ligand binding relative to the nucleotide free or MgADP bound states. In line with these previously reported iron chelation studies, Se-incorporation into the Fe_4_S_4_ cluster is only observed in the presence of MgATP. The solvent accessibility of the Fe protein cluster^17,29–31^ contrasts with most Fe_4_S_4_ containing proteins that feature buried clusters, with only a few exceptions.^30,32^ It should be noted that while the Fe protein cluster remains relatively exposed in the absence of nucleotide or in the presence of MgADP (Figure 2a and 2b), incubation with KSeCN does not result in chalcogenide exchange under these conditions (Figure 2-figure supplement 2d and 2e). Consequently, the position of the cluster near the surface of the protein is not a sufficient condition for KSeCN-derived Se-incorporation. These observations highlight how the MgATP-dependent reactivity of the Fe protein cluster represents an element of control, both for the physiological properties as well as cluster atom exchange.

While the crystallographic observations described herein unambiguously establish the occurrence of chalcogenide exchange at the Fe protein cluster, the mechanism of this reaction remains open. The ability of Fe protein to reduce CO_2_-to-CO, in the absence of the MoFe protein, suggests that the Fe_4_S_4_ cluster may coordinate CO_2_.^4^ Furthermore, the first observed instance of N_2_ bound to a synthetic cluster (a MoFe_3_S_4_ cubane) was recently reported, demonstrating that relatively simple FeS clusters can coordinate ligands.^33^ In the context of MoFe protein-independent CO_2_ reduction and N_2_-binding to a synthetic cluster, KSeCN can be viewed as a substrate analogue to CO_2_, with the Se-exchange mechanism proceeding by initial ^-^SeCN binding to an Fe center, followed by Se-C bond cleavage, and chalcogenide exchange. The exchange of selenium into an Fe_4_S_4_ cluster through reaction with KSeCN represents a new approach for generating Fe_4_Se_4_-containing Fe proteins from native Fe_4_S_4_-containing proteins and a selenium source that complements existing reconstitution approaches using apoproteins (proteins deficient in the native Fe_4_S_4_ cluster) and a (i) selenium source, iron source, and reductant or (ii) with synthetic clusters.^34,35^ The capability of preparing chalcogenide substituted clusters opens new avenues for exploring the multiple roles of the Fe protein in biological nitrogen fixation.

## Methods

### General considerations

All protein manipulations were carried out using standard Schlenk or anaerobic tent techniques under an atmosphere of Ar or 97/3% Ar/H_2_ mixture, respectively. Potassium selenocyanate (KSeCN) was purchased from Sigma Aldrich. All other reagents were purchased from commercial vendors and used without further purification unless otherwise stated.

### Growth of *A. vinelandii* and nitrogenase purification

*Azotobacter vinelandii* Lipman (ATCC 13705, strain designation OP) growth and nitrogenase purification were performed based on previously published methods^36,37^ with the following modifications. All protein buffers (pH 7.8) were deoxygenated, kept under an argon atmosphere, and contained 5 mM dithionite (Na_2_S_2_O_4_). The supernatant from the centrifuged cell lysate was loaded onto a Q Sepharose fast flow column (GE Healthcare). *In vitro* nitrogenase activity was determined by monitoring acetylene reduction to ethylene as previously described.^14^ Ethylene and acetylene were quantified using gas chromatography (activated alumina 60/90 mesh column, flame ionization detector). MoFe protein had a specific activity of 2940 ± 30 nmol min^-1^ mg^-1^ (V_max_) and Fe protein had a specific activity of 1880 ± 90 nmol min^-1^ mg^-1^ (V_max_) when measured by acetylene reduction at saturation of each component.

### Preparation of Se-Incorporated Nitrogenase Proteins Using KseCN

The Se-incorporated proteins were prepared using a previously reported protocol,^14^ with the following modifications. To generate sufficient material for EPR spectroscopy or crystallization, two parallel 12 mL reactions (each containing 1.5 mg of MoFe protein and 1.65 mg of Fe protein [component ratio of 2]) were combined and concentrated under argon overpressure using an Amicon filtration cell with a molecular weight cut-off of 100 kDa. The resultant concentrated protein was used to crystallize Se-incorporated MoFe protein. The corresponding 100 kDa *filtrate* was collected, and re-subjected to concentration under argon overpressure using an Amicon filtration cell with a molecular weight cut-off of 30 kDa. The latter batch of concentrated protein was used to crystallize Se-incorporated Fe protein. Note that the filter membranes did not completely separate the Se-incorporated proteins (as determined by SDS-PAGE); regardless, selective crystallization of either protein was successful (*vide infra*).

### Control KSeCN reactions with no MoFe protein

The procedure for the various control reactions were identical to that of the preparation of Se-incorporated nitrogenase proteins described above with the following changes noted. No MoFe protein was included in the control reactions. Because the MoFe protein was absent in these reactions, a 30 kDa filter membrane was used to concentrate the reaction mixture for crystallization. In addition, for the no-nucleotide control, the components of the ATP regeneration system were excluded and the resultant concentrated protein was rinsed with a 5 mM MgADP solution (3 × 8 mL) for crystallization purposes. Finally, for the MgADP control, the ATP regeneration system was replaced with a 5 mM MgADP solution.

### Crystallization and data collection of Se-incorporated MoFe protein

The Se-incorporated MoFe protein was crystallized by the sitting-drop vapor diffusion method at ambient temperature in an inert gas chamber. The reservoir solution contained 15-20% polyethylene glycol (PEG) 4000, 0.5-0.8 M NaCl, 0.2 M imidazole/malate (pH 8.0), and 5 mM dithionite. Additionally, native MoFe protein crystals (crushed using a seed bead Eppendorf tool with either a plastic bead or glass beads) were used as seeds to accelerate the crystallization process and improve the overall crystal quality. For flash-cooling, 2-methyl-2,4-pentanediol (MPD) was either added directly to the crystal droplet, yielding 10% MPD, or the crystals were transferred into a harvesting solution consisting of the reservoir solution and 10% MPD. Complete sets of diffraction data were collected at the Synchrotron Radiation Lightsource (SSRL) beamline 12-2 equipped with a Dectris Pilatus 6M detector. Two sets of anomalous diffraction data were collected above and below the Se K-edge at 12,668 eV (0.978690 Å) and 12,643 eV (0.980620 Å), respectively. Data were indexed, integrated, and scaled using iMosflm, XDS, and Aimless.^38–40^ Phase information were obtained using the available 1.00 Å resolution structure (PDB: 3U7Q) as a molecular replacement model, omitting the metalloclusters and water from 3U7Q. Structural refinement, and rebuilding were accomplished by using REFMAC5/ PHENIX, and COOT, respectively.^41–43^ Neutral atomic scattering factors were used in the refinement. Anomalous difference Fourier maps were calculated using CAD/FFT in the CCP4 suite. The double difference anomalous Fourier maps were calculated using SFTOOLS (CCP4). Protein structures were displayed in PYMOL.

Consistent with our previously published MoFe protein structures containing Se-incorporated FeMo-cofactor,^14,44^ this structure revealed that (1) the belt sulfides were labile, with Se-incorporation predominantly at the 2B site, but also at the 5A and 3A sites (Figure 2-figure supplement 5) and (2) no Se-incorporation occurs at the P-cluster.

### Preparation, crystallization and data collection of Se-incorporated Fe protein

Se-incorporated Fe protein was crystallized by the sitting-drop vapor diffusion method at ambient temperature in an inert gas chamber. The reservoir solution contained 36-41% PEG 400, 0.1-0.3 M NaCl, 0.1 M 4-(2-hydroxyethyl)-1-piperazineethanesulfonic acid (HEPES) (pH 7.5), 2.5 mM dithionite, and 0.17 mM 7-cyclohexyl-1-heptyl-ß-D-maltoside (Cymal 7). The same parameters for data collection and refinement as Se-incorporated MoFe protein were used, with the following modifications: phase information was obtained using PDB coordinate set 6N4L as the Fe protein molecular replacement model, with the cluster, MgADP, and water molecules omitted. Cluster modeling was accomplished by modeling individual X (X = Se, S) and Fe atoms at the respective cluster positions and by inputting bond distance and bond angle restraints, based on the core cluster metrics determined for synthetic clusters (SIMNOR10 and COZXUK), into the PHENIX.REFINE configuration.^45,46^ The *f’* = −6.00 and *f”* = 4.00 values for Se were used, with the latter value matching well with the fluorescence scans of Se-incorporated Fe protein crystals (see Figure 2-figure supplement 6 for sample fluorescence scan). Se occupancies were determined by fixing the cluster atom B-factors to the value the Fe atoms refined to during an initial refinement. Given that B-factors and occupancies are correlated and the fact that there is minimal difference between the S and Fe cluster atom B-factors in Se-free crystals (see Figure 2-figure supplement 2 and Supplementary file 2), this approach is reasonable. Neutral atomic scattering factors were used in the refinement. Anomalous difference Fourier maps were calculated using CAD/FFT in the CCP4 suite. The double difference anomalous Fourier maps were calculated using SFTOOLS (CCP4). Protein structures were displayed in PYMOL. Given restrictions regarding cluster notation as determined by the PDB, the individual atom notation in our models was converted to the cluster (SFS or SF4) format for the purposes of depositing the structures into the PDB. While the two-cluster model accurately reflects the occupancies at the distinct chalcogenide sites (X3 and X4) determined upon refinement with the individual atom cluster notation, we recognize that the two-cluster model does not realistically reflect the data and that a mixture of partially occupied Se-incorporated clusters is likely, i.e. Fe_4_S_4_, Fe_4_S_3_Se, Fe_4_S_2_Se_2_, Fe_4_SSe_3_, and Fe_4_Se_4_ may all be present to yield the crystallographically determined occupancies.

The structural models and structure factors have been deposited with the Protein Data Bank (PDB) under accession codes 7TPW, 7TPX, 7TPY, 7TPZ, 7T4H, 7TQ0, 7TQ9, 7TQC, 7TNE, 7TQE, 7TQF, 7TPN, 7TQH, 7TQI, 7TPO, 7TQJ, 7TQK, and 7TPV. For tables with data collection and refinement statistics, please see Supplementary files 4-9.

### KSeCN-soaking of Fe protein crystals

The MgADP bound crystal form of the Fe protein was soaked with KSeCN (5 mM) by adding KSeCN directly to a crystal well, re-sealing, and allowing the well to sit for various lengths of time. The particular dataset provided here was obtained after the crystals had been soaked with KSeCN for one week.

### Purified Se-incorporated Fe protein EPR sample preparation

Se-labeled protein from three KSeCN reaction sets were combined and loaded onto an anaerobic 1 mL HiTrap Q anion exchange column (previously equilibrated with 50 mM tris/HCl buffer (pH = 7.8) which contained 150 mM NaCl [low salt] and 5 mM dithionite). Se-incorporated MoFe protein and Se-incorporated Fe protein eluted with a linear NaCl gradient at 280 and 430 mM NaCl, respectively. Se-incorporated Fe protein was concentrated to approximately 16 mg/ml under argon overpressure using an Amicon filtration cell with a molecular weight cut-off of 30 kDa. The EPR sample was prepared as an approximately 50 μM frozen glass of Se-incorporated Fe protein in a 50:50 mixture of buffer:ethylene glycol. The buffer solution consisted of 200 mM NaCl and 50 mM tris/HCl (pH = 7.8) and contained 25 mM dithionite (7.5 mM dithionite in EPR sample overall).

### CW EPR Spectroscopy

X-band EPR spectra were obtained on a Bruker EMX spectrometer equipped with an ER 4116DM Dual Mode resonator operated in perpendicular mode at 10 K using an Oxford Instruments ESR900 helium flow cryostat. Bruker Win-EPR software (ver. 3.0) was used for data acquisition. Spectra were simulated using the EasySpin^47^ simulation toolbox (release 5.2.28) with Matlab 2020b.

### Discussion of EPR data

The Fe protein features an *S* = 1/2 signal corresponding to the [Fe_4_S_4_]^1+^ state of the cluster with *g* = [2.05, 1.94, 1.88].^27^ While the Fe protein can exist in the *S* = 3/2 and *S* = 1/2 states, the population of the spin state depends on the sample conditions, including the presence of nucleotide and solvent. In 50% ethylene glycol, used as a cryoprotectant, most of the Fe protein cluster is in the *S* = 1/2 state.^48^

The mixture of S/Se-labeled Fe protein could be separated from the MoFe protein using anion exchange chromatography and subjected to electron paramagnetic resonance (EPR) spectroscopy. Based on the crystallographic data, we anticipate that the Se-labeled Fe protein exists in a mixture of Se-containing cluster states (i.e. Fe_4_S_4_, Fe_4_S_3_Se, Fe_4_S_2_Se_2_, Fe_4_SSe_3_, and Fe_4_Se_4_ may all be present). As such, a familiar *g* = 2 signal corresponding to the [Fe_4_S_4_]^1+^ cluster of the Fe protein was observed (Figure 2-figure supplement 7). While there are slight differences in the EPR spectra between the all-S vs Fe_4_X_4_ (X = S, Se) mixture of the Fe protein cluster, the signal of the -S/-Se mixture could be successfully simulated using the same parameters as the all-S containing Fe protein cluster.^49^ One plausible interpretation of our EPR data is that the various Fe_4_X_4_ states yield nearly identical, overlapping, signals consistent with the observation that EPR spectra of Fe_4_S_4_ vs Fe_4_Se_4_ clusters are nearly identical.^50^ Alternatively, it has been recently reported that an Fe protein with an Fe_4_Se_4_ cluster is reduced to the all ferrous state in the presence of dithionite, rendering it EPR silent in perpendicular mode EPR.^35^ In this context, the signal observed in Figure 2-figure supplement 7 may correspond to the [Fe_4_S_4_]^1+^ state while the [Fe_4_Se_4_]^0^ state is not observed. Our results cannot distinguish between these two possible interpretations.

## AUTHOR INFORMATION

### Notes

The authors declare no competing financial interests.

## ACKNOWLEDGMENT

*We dedicate this paper to Prof. James B. Howard and are grateful for our extensive exchanges on cluster atom exchanges—Jim really shaped the way we thought about the Fe protein cluster and our journeys with nitrogenase*. We also thank Dr. Javier Fajardo Jr. for insightful discussions, Jeffrey Lai for growing *Azotobacter vinelandii*, and Dr. Paul Oyala for EPR training and support. The authors are grateful to the Gordon and Betty Moore Foundation, Don and Judy Voet, and the Beckman Institute at Caltech for their generous support of the Molecular Observatory at Caltech. Use of the Stanford Synchrotron Radiation Lightsource, SLAC National Accelerator Laboratory, is supported by the U.S. Department of Energy, Office of Science, Office of Basic Energy Sciences under Contract No. DE-AC02-76SF00515. The SSRL Structural Molecular Biology Program is supported by the DOE Office of Biological and Environmental Research, and by the National Institutes of Health, National Institute of General Medical Sciences (including P41GM103393). The Caltech EPR Facility is supported by NSF-1531940. This work was supported by the National Institute of Health (NIH Grant GM45162) and the Howard Hughes Medical Institute (HHMI). This article is subject to HHMI’s Open Access to Publications policy. HHMI lab heads have previously granted a nonexclusive CC BY 4.0 license to the public and a sublicensable license to HHMI in their research articles. Pursuant to those licenses, the author-accepted manuscript of this article can be made freely available under a CC BY 4.0 license immediately upon publication.

---

**Figure 2- figure supplement 1.**
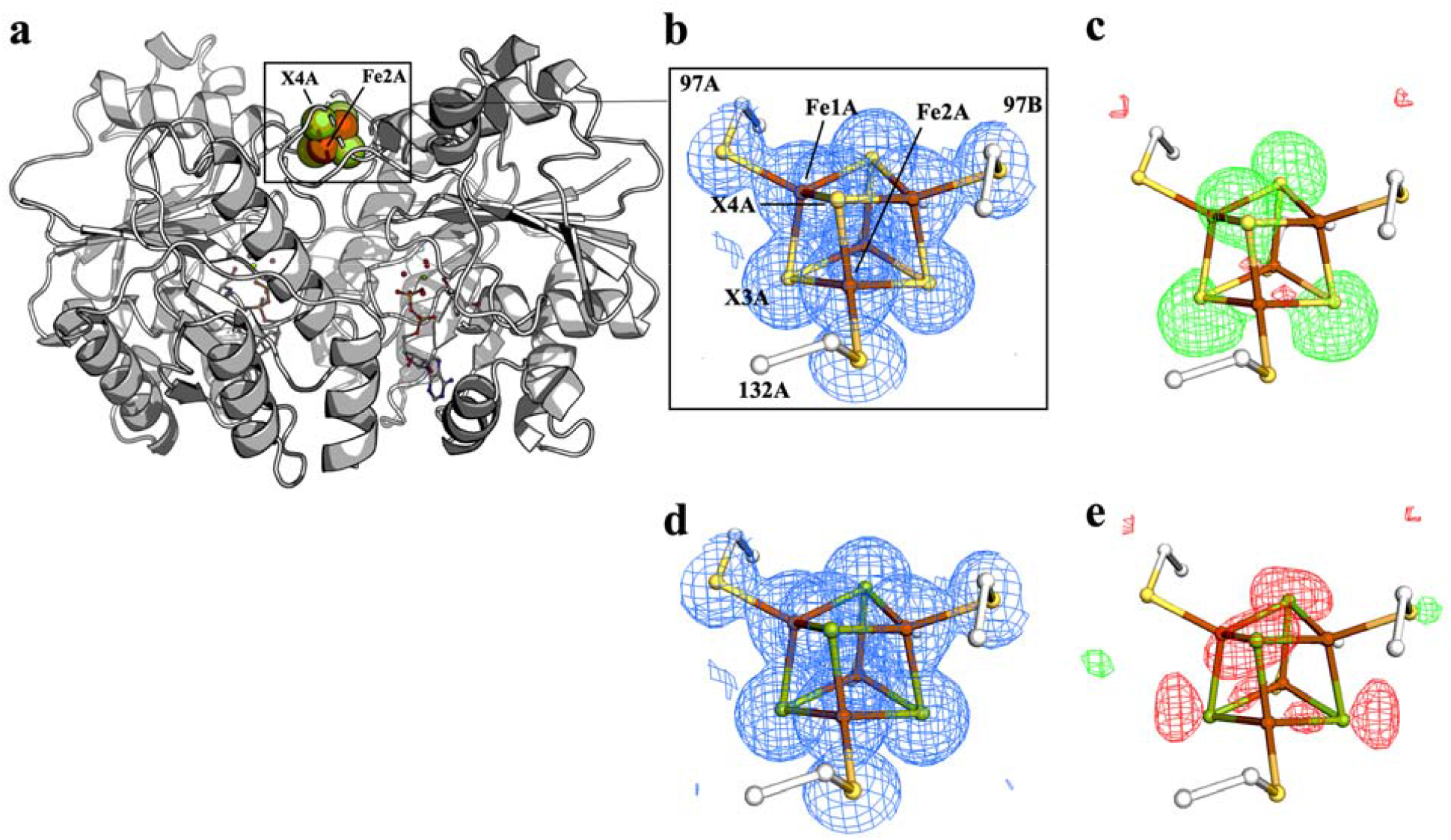
Comparison of 2F_obs_-F_calc_ and F_obs_-F_calc_ maps for cluster modeled as exclusively S- vs Se-containing forms. (a) Protein overview. The all S-containing model with (b) overlaid electron density (2F_obs_-F_calc_) map contoured at 1.5 σ (blue mesh) and (c) overlaid difference density (F_obs_-F_calc_) map contoured at 3.0 σ. The all Se-containing model with (d) overlaid electron density (2F_obs_-F_calc_) map contoured at 1.5 σ (blue mesh) and (e) overlaid difference density (F_obs_-F_calc_) map contoured at 3.0 σ. The positive and negative difference densities in (c) and (e) are displayed in green and red, respectively.

**Figure 2- figure supplement 2.**
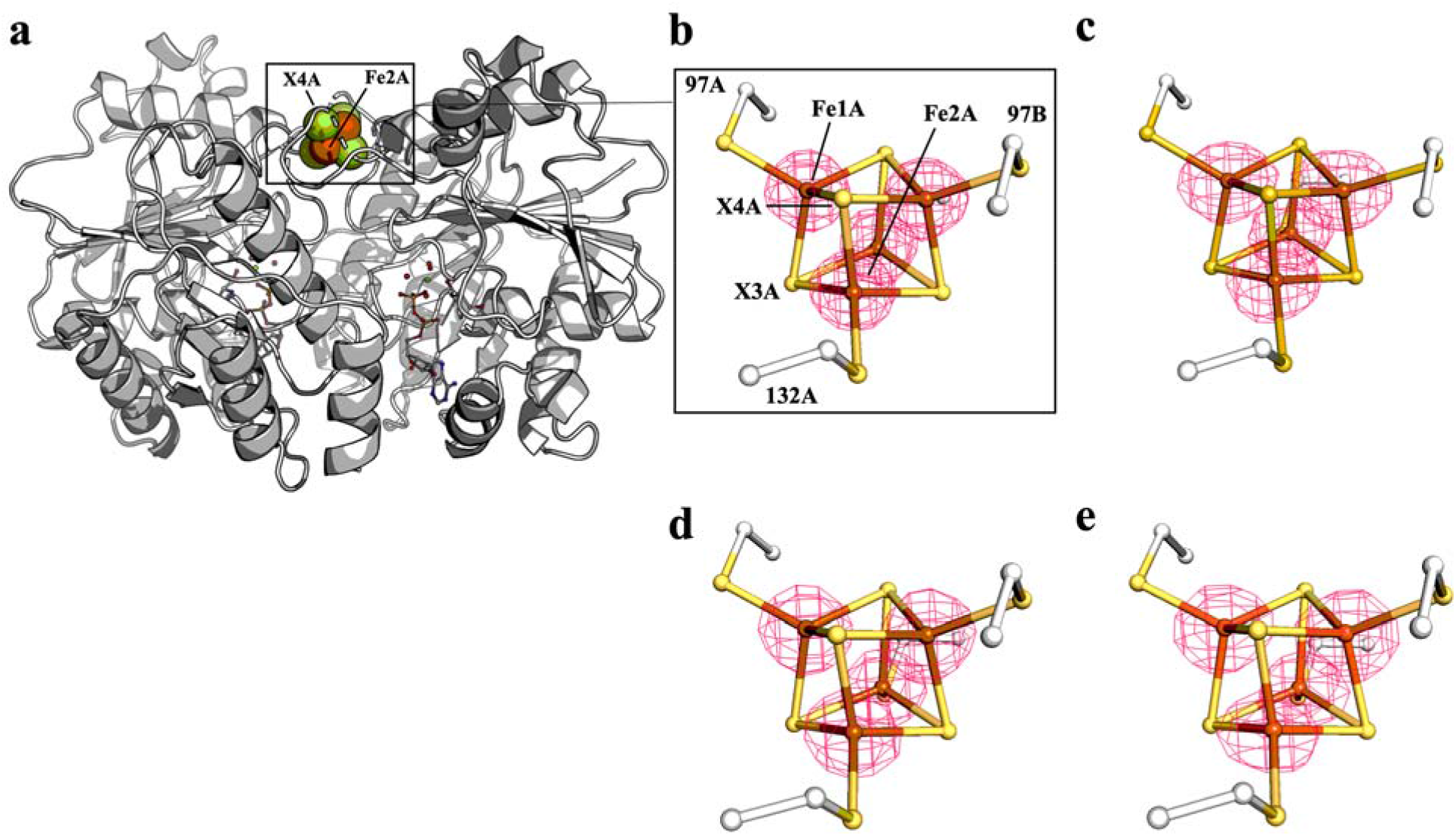
Anomalous difference Fourier maps (pink mesh, contoured at 7.0 σ) calculated from diffraction data collected at 12,668 eV for Se-free Fe protein crystals corresponding to crystal in (b) Supplementary file 2, PDB ID 7TPW (c) Supplementary file 2, PDB ID 7TPX, (d) Supplementary file 2, PDB ID 7TPY (nucleotide free reaction), and (e) Supplementary file 2, PDB ID 7TPZ (MgADP in place of MgATP/ATP regeneration system). A protein overview is shown in (a) for orientation purposes.

**Figure 2- figure supplement 3.**
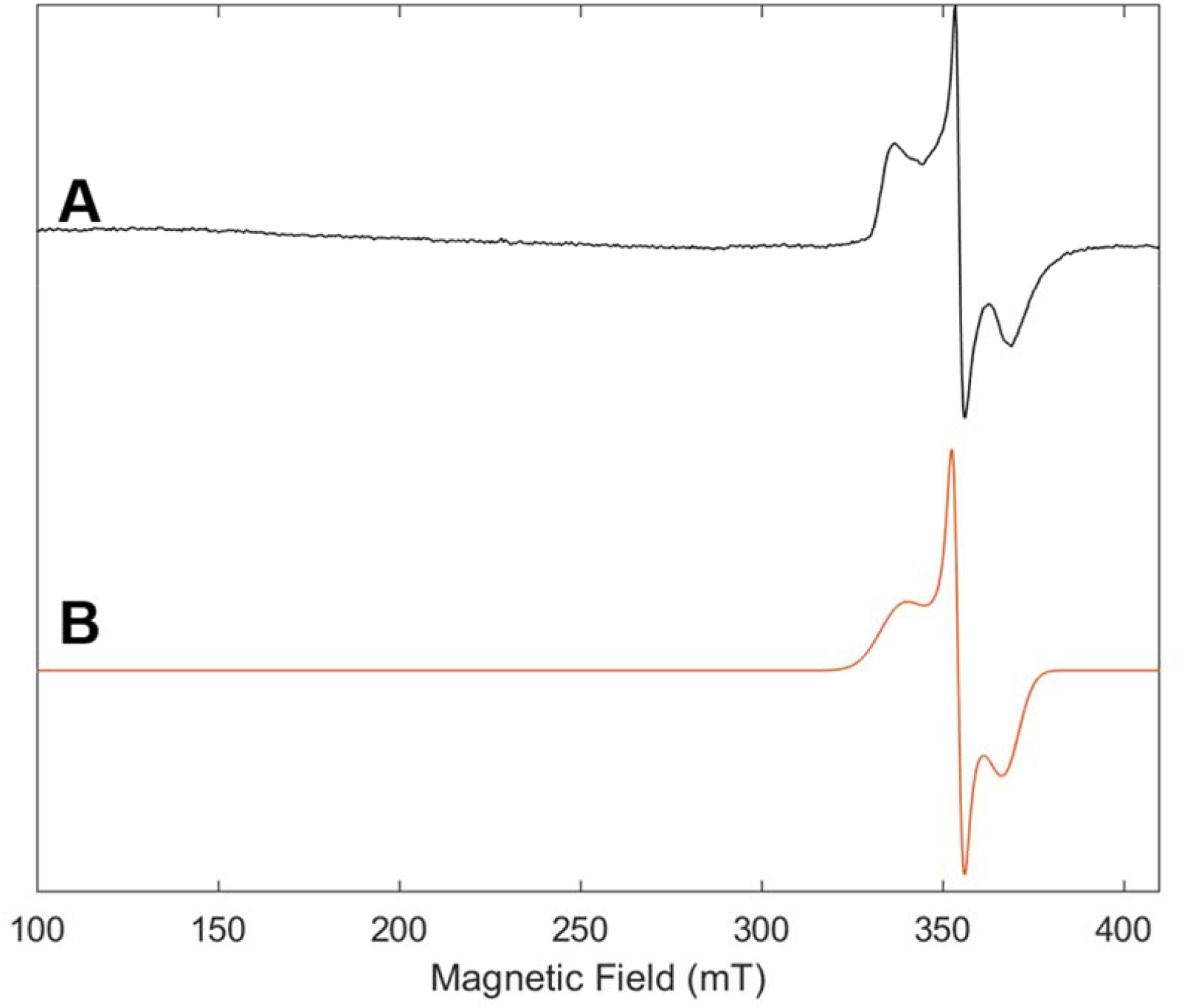
EPR spectrum of purified Fe protein used in control KSeCN reaction with no MoFe protein with **(A)** experimental data (black) and **(B)** simulation (orange). The frozen solution (10K) spectrum was collected at 9.64 GHz with a microwave power of 2.2 mW, a modulation amplitude of 8.0 G, a modulation frequency of 100 KHz and conversion time of 42 ms.

**Figure 2- figure supplement 4.**
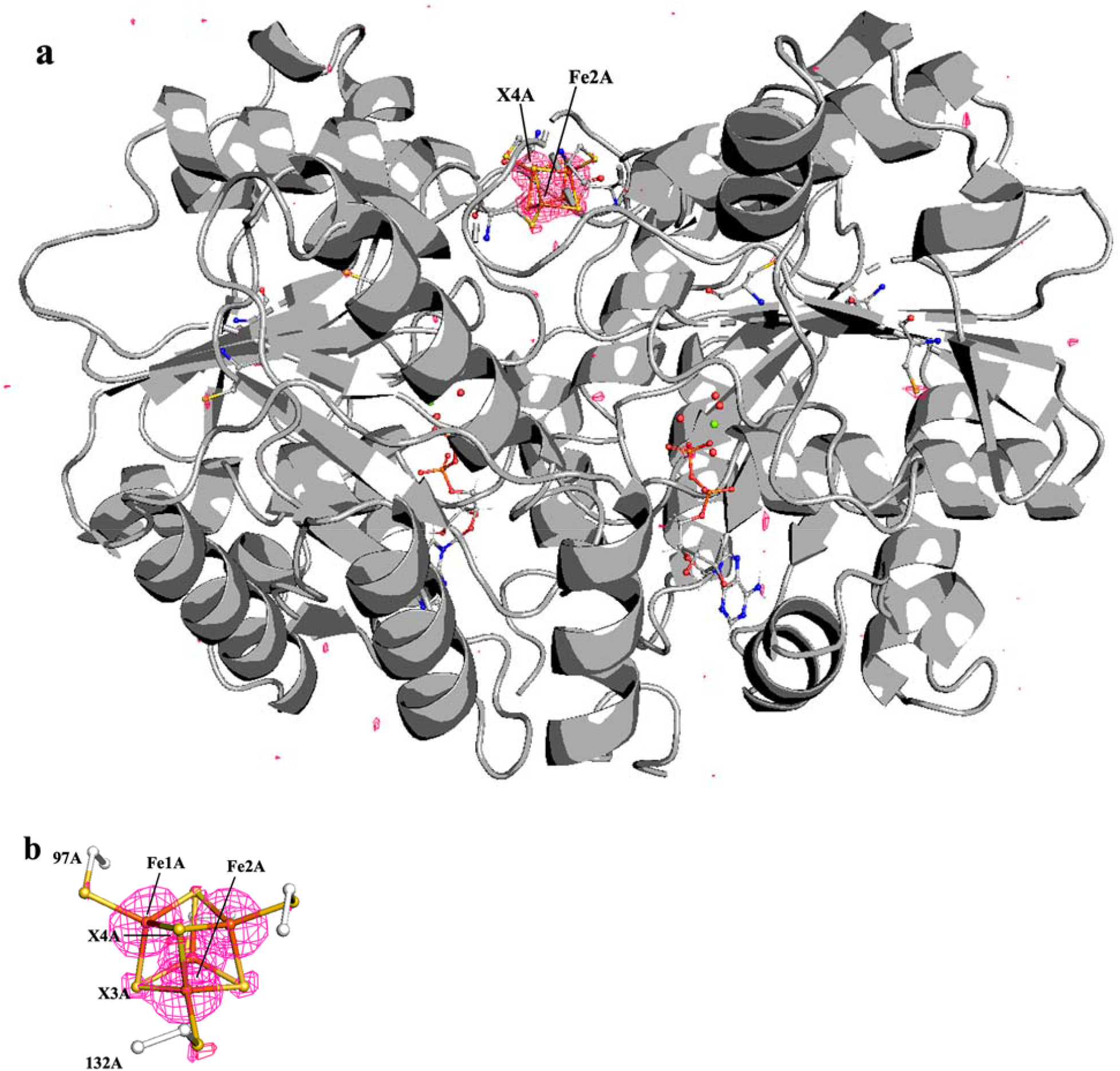
Anomalous difference Fourier map (pink mesh, contoured at 4.0 σ) calculated from diffraction data collected at 12,668 eV for Fe protein crystal soaked with KSeCN shown over (a) the entire protein, (b) the cluster.

**Figure 2- figure supplement 5.**
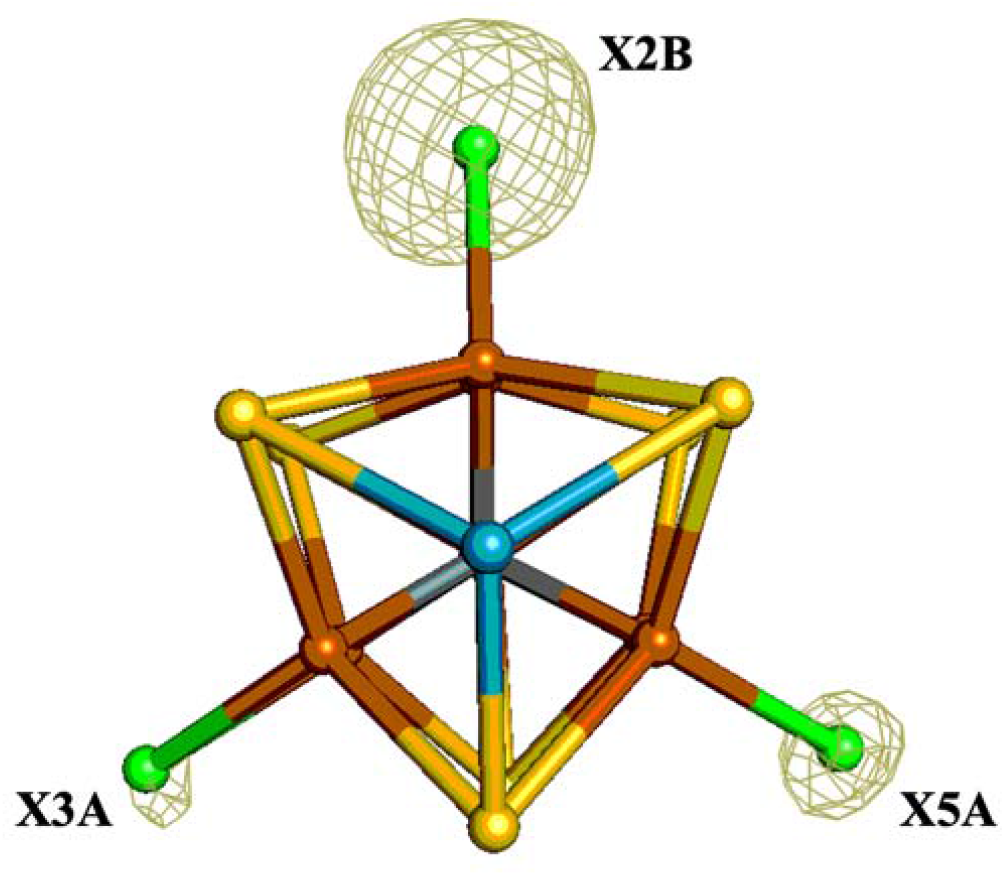
Pymol representation of the FeMo-cofactor of Se-incorporated MoFe protein overlaid with the double difference (Δanom_12668eV_–Δanom_12643eV_) anomalous map contoured at 9.0 σ (olive mesh); X denotes a mixture of Se and S atoms. Iron atoms are shown in orange, sulfur in yellow, molybdenum in turquoise, and a mixture of Se/S in green.

**Figure 2- figure supplement 6.**
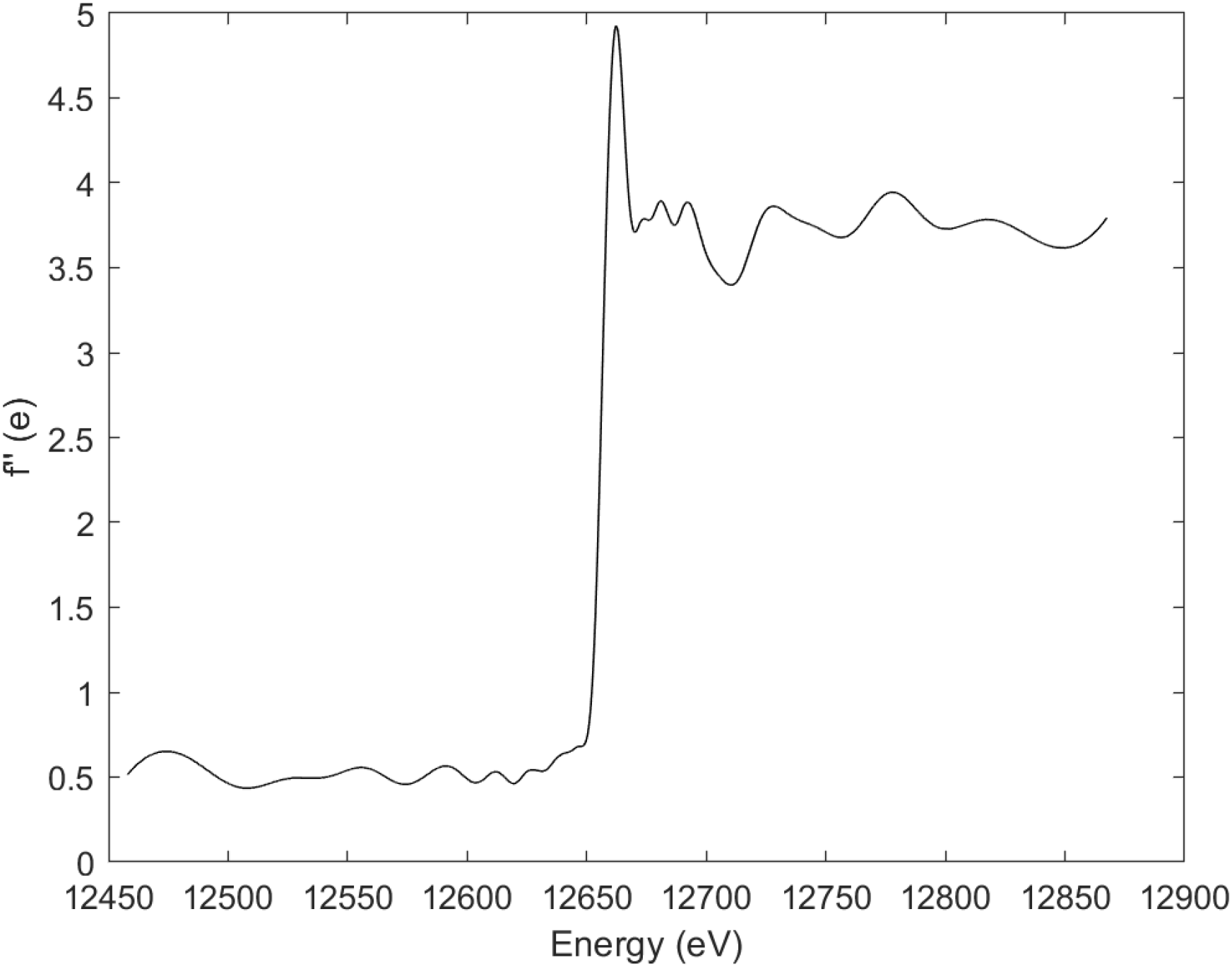
Fluorescence scan collected around Se K-edge (12658 eV) for Se-incorporated Fe protein.

**Figure 2- figure supplement 7.**
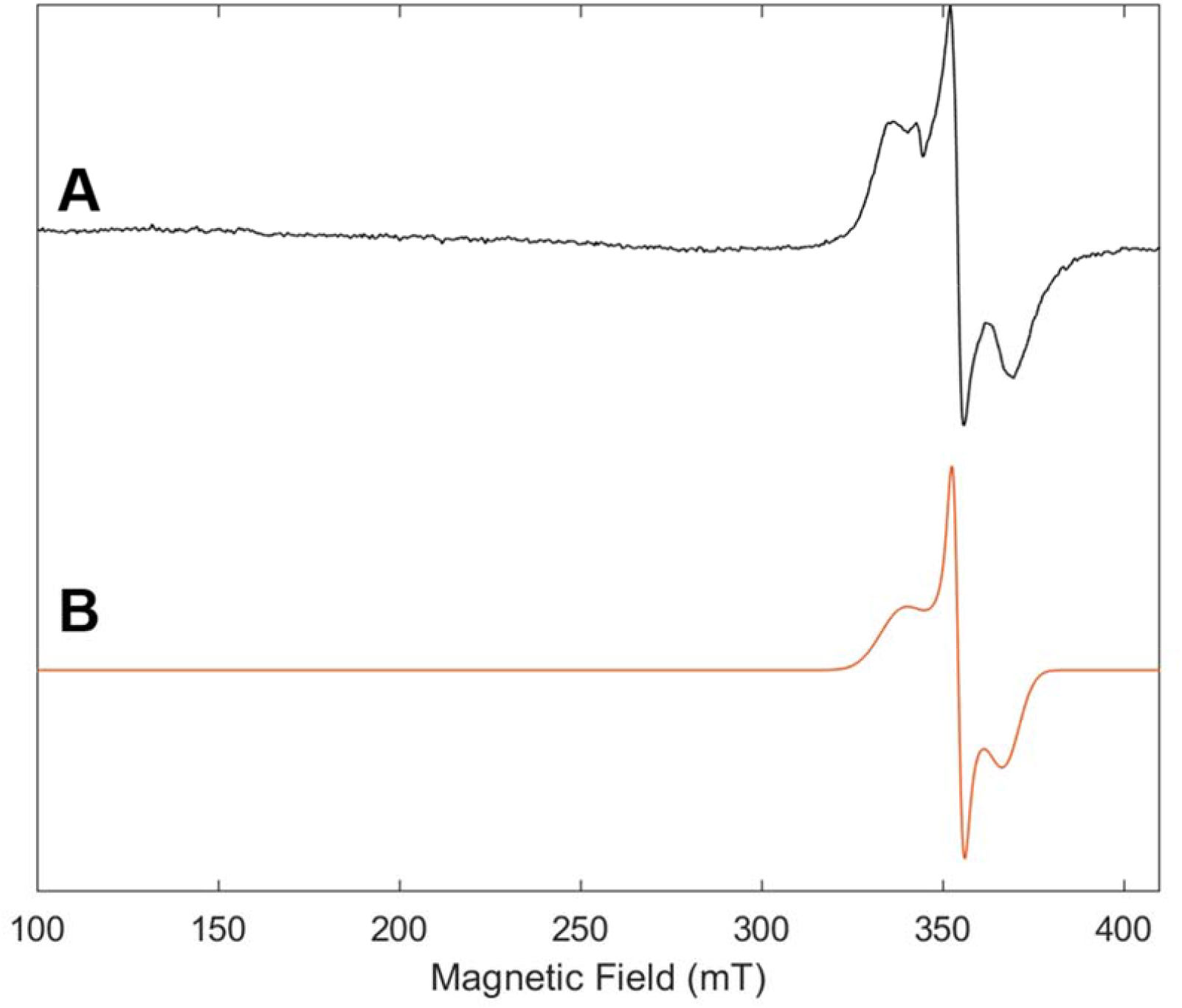
EPR spectrum of purified Se-incorporated Fe protein with **(A)** experimental data (black) and **(B)** simulation (orange). The frozen solution (10K) spectrum was collected at 9.64 GHz with a microwave power of 2.2 mW, a modulation amplitude of 8.0 G, a modulation frequency of 100 KHz and conversion time of 42 ms. For EPR spectrum of Fe protein, see Figure 2- figure supplement 3.

